# Ultrastructure of the brain waste-clearance pathway

**DOI:** 10.1101/2023.04.14.536712

**Authors:** Søren Grubb

**Affiliations:** Center for Translational Neuroscience and Department of Neuroscience, Faculty of Health Sciences, University of Copenhagen, DK-2200 Copenhagen N, Denmark

**Keywords:** macrophage, fibroblast, leptomeninges, metabolic waste, GFAP

## Abstract

Efficient metabolic waste clearance is essential for optimal brain function. Waste accumulation can cause inflammation, tissue damage, and potentially lead to dementia and other neurological disorders. The primary waste exit route is debated, with evidence suggesting the lymphatic system or arachnoid villi/granulations as possible pathways.

This study uses an ultrastructural dataset to emphasize the potential underestimation of macrophages and arachnoid villi/granulations in waste removal.

Based on brain surface ultrastructure analysis of the MICrONS dataset, waste clearance may occur through a transcellular route involving astrocytes to the perivascular space. Waste is then either phagocytosed by perivascular macrophages or directed to the subarachnoid space for meningeal macrophage phagocytosis.

Evidence is provided that macrophages transport non-degradable waste in lysosomes to arachnoid villi for exocytosis into the villus lumen and venous system routing. Additionally, the study investigates fibroblast-macrophage interactions via primary cilia, potentially optimizing waste clearance. The role of astrocytes in brain water homeostasis and GFAP’s potential significance in water filtration are also examined. This study offers valuable insights into waste clearance mechanisms in the brain, potentially aiding dementia understanding and treatment.

## Introduction

Accumulation of brain metabolic waste can lead to inflammation and tissue damage, potentially causing dementia and other neurological disorders. While the current focus of brain-waste clearance is on the flow of cerebrospinal fluid (CSF) out of the brain via a lymphatic route(Rasmussen, Mestre, and Nedergaard 2022), recent research has highlighted the role of macrophages and microglia in waste removal(Drieu et al. 2022; El Hajj et al. 2019; Karam et al. 2022; Kierdorf et al. 2019; Spangenberg et al. 2019; Wendt et al. 2022). Macrophages and microglia are immune cells responsible for engulfing and eliminating cellular debris, pathogens, and other harmful substances, including metabolic waste. In this study, I use an ultrastructural dataset to explore the mechanisms behind brain macrophage phagocytosis and waste disposal, as well as their potential implications for brain health and disease.

The glymphatic system, which involves the perivascular pumping of CSF, also plays an important role in metabolic waste clearance(Rasmussen et al. 2022). The neurovascular unit (NVU) is a key player in this system(Bojarskaite et al. 2022; Fultz et al. 2019; Grubb 2022; Grubb and Lauritzen 2019; Grubb, Lauritzen, and Aalkjær 2021), but its exact involvement is still unclear. Moreover, functional hyperemia, which increases blood flow to meet neuronal demands, may have an additional purpose, such as aiding in the removal of metabolic waste.

The arachnoid villi and granulations, which are protrusions of the arachnoid mater that reach the dural veins / venous sinuses, have long been thought to be a route for CSF outflow from the brain. However, recent research suggests that lymph vessels may be the primary route for CSF exit(Ma et al. 2017; Proulx 2021). The filtration mechanism of arachnoid villi and granulations also remains elusive, and it is unclear whether they play a role in CSF reabsorption.

In addition, I will discuss the use of PEGylated tracers, which can bypass macrophages, and the potential for bias in using tracers that may end up in the lymphatic route due to high concentrations. It is important to note that this study is purely observational and is intended to generate hypotheses that can be tested in future research.

Overall, these findings may have broad implications for the study of diseases like dementia, subarachnoid hemorrhage, and meningitis.

## Results

The MICrONS dataset includes the surface of the brain, specifically the pia- and arachnoid mater (collectively known as leptomeninges), but these structures have not been segmented (Figure 1a). Recently, the leptomeninges have attracted significant interest due to their potential role in removing cerebrospinal fluid (CSF) and metabolic waste from the brain (Møllgård et al. 2023). To segment these structures, I annotated tissue, cell, and organelle ultrastructures using Neuroglancer, a web-based viewer, and generated 3D renderings of these structures. Initially, I present the structures of the glia limitans, pial blood vessels, arachnoid mater, and Virchow-Robin space (Figure 1b and c, and Video 1). Subsequently, I discuss the ultrastructure of the cells found in these regions. The isolated glia limitans appears furrowed, indicating the locations of large pial arterioles and venules, which are covered by the arachnoid mater (Figure 1b).

**Figure 1.**
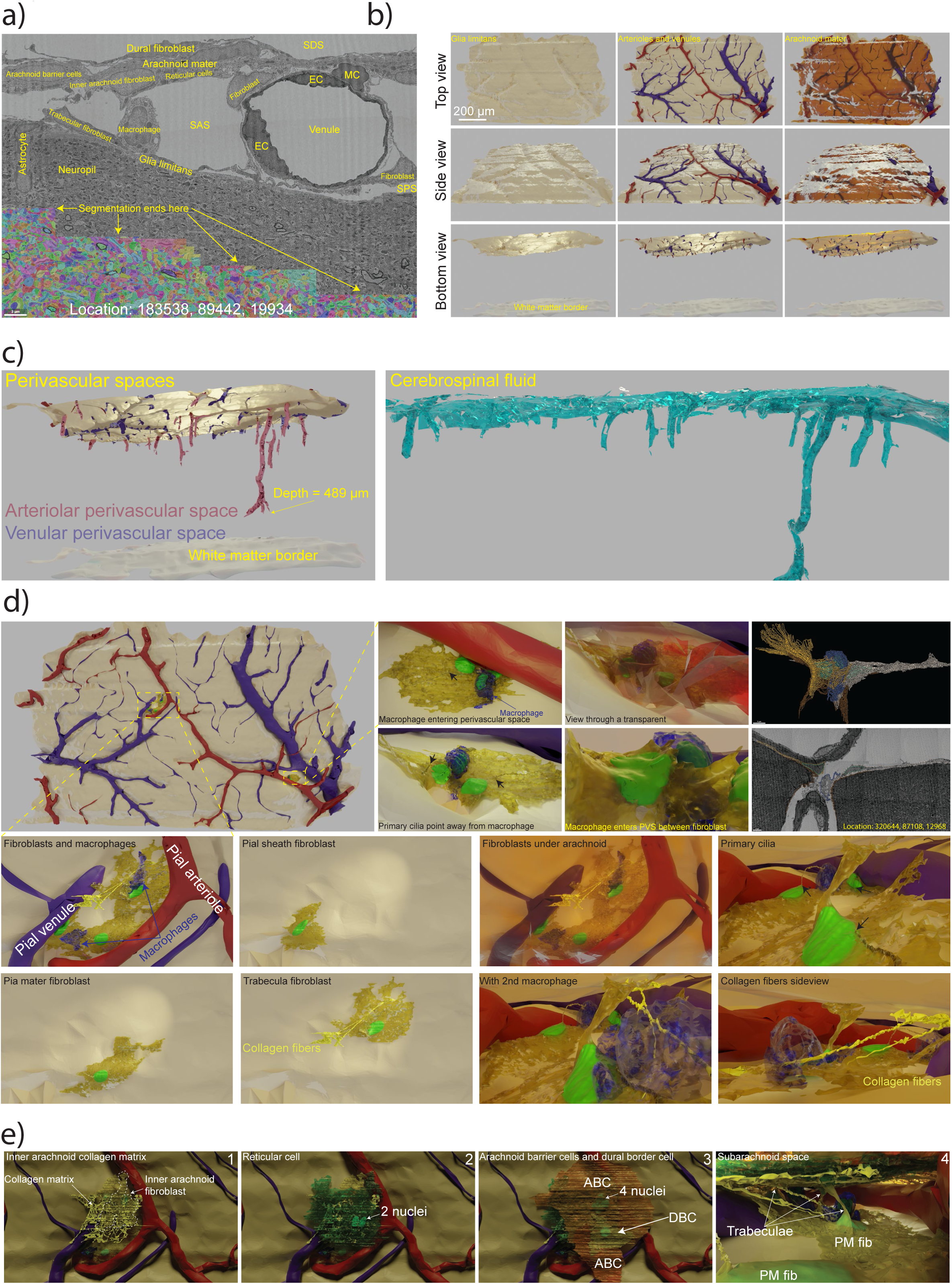
Perivascular spaces and fibroblasts. **a)** The MICrONS dataset segmentation border extends approximately 10 µm below the glia limitans. The leptomeninges consist of three spaces: the subdural space (SDS), the subarachnoid space (SAS), and the subpial space (SPS). A cross-section of a pial venule within the SAS displays annotated typical cells: MC = mural cell, EC = endothelial cell. **b)** Rendered brain surface shown from three different angles, highlighting the outer limit of the glia limitans, pial vasculature lumen, and inner limit of the inner arachnoid fibroblasts. Notice how blood vessels correspond to furrows in the glia limitans. The outer limit of the white matter illustrates the cortex’s reduced size. **c)** *Left panel:* PVSs in pink for pial arteries (PAs) and light purple for ascending venules, illustrating the difference in PVS depths. *Right panel:* CSF is illustrated by combining the outer limit of the glia limitans with the inner limit of the inner arachnoid fibroblasts and PVSs, while subtracting the vascular lumen. **d)** *Upper left panel:* Overview of two locations with annotated and rendered fibroblasts. *Upper right panels:* Macrophage entering the PVS of a PA between processes of two pia mater fibroblasts. Black arrows indicate primary cilia of fibroblasts. The furthest right panels display annotation data in 3D and ultrastructure annotations. *Lower panels:* Three pia mater fibroblasts and two macrophages rendered in a space between a pial arteriole and venule. One fibroblast serves as a pure pia mater fibroblast, another partially forms the venule’s pial sheath, and the third fibroblast forms an arachnoid trabecula. Collagen fibers (yellow) extend from the arachnoid mater to the subpial space. The lower right panels depict three different views with the arachnoid mater included. Notice how all three fibroblasts have primary cilia in contact with macrophages. **e)** Segmentation of cells and structures constituting the arachnoid mater at the same location. 1) Inner arachnoid fibroblast (golden with white dashed outline) beneath a collagen matrix (yellow) in the inner arachnoid. 2) Large reticular cell with 2 nuclei (green). 3) A large and a small arachnoid barrier cell (brown) with 4 and 1 nuclei respectively. Dural border cell (golden) on top of where the two arachnoid barrier cells meet. 4) A view from the subarachnoid space with the pia mater and inner arachnoid trabeculae reaching across the SAS.

### Perivascular spaces (Virchow-Robin spaces)

Virchow-Robin spaces, which surround brain parenchymal blood vessels, have been referred to as either perivascular or paravascular spaces(Bacyinski et al. 2017; Wardlaw et al. 2020).This distinction arises from the presence of a meningeal fibroblast layer, known as the pial sheath, which surrounds the pial and penetrating vessels(Zhang, Inmant, and Wellert 1990). Perivascular spaces are believed to contain interstitial fluid, as they are bounded internally by the vascular basement membrane and externally by either fibroblasts or, in the absence of fibroblast coverage, by the glia limitans. Paravascular spaces, on the other hand, contain cerebrospinal fluid (CSF) since they are continuous with the subarachnoid space and are internally bounded by pial sheath fibroblasts and externally by the glia limitans.

Due to the non-continuous nature of the pial sheath, particularly after arterioles penetrate the parenchyma (Zhang et al. 1990), the distinction between perivascular and paravascular spaces is ambiguous. Consequently, the more widely used term "perivascular spaces" (PVS) has been adopted for Virchow-Robin spaces in this study. When discussing spaces around pial vasculature, I will use the term "subarachnoid space" (SAS).

In the MICrONS dataset, PVSs can be found surrounding 21 penetrating arterioles (PAs) at depths ranging from 25-489 µm below the brain surface (Figure 1c, Sup. Table 1, Segmentations: tinyurl.com/2323bdwx). The average depth of the arteriolar PVSs in this dataset is 125±22µm. No PVS exists around ascending venules beyond the first 15-20µm below the brain surface, consistent with MRI studies(Wardlaw et al. 2020). As previously described, in cases where PVSs of penetrating arterioles (PA) extend to capillaries, they terminate at the 1^st^-order capillary(Grubb 2022). For one PA, the PVS reaches 489 µm below the pial surface (Figure 1c, Location: 320752, 84727, 12738); however, the first branch point of that PA is at 423 µm. The PVSs not only contain CSF but also metabolic waste products, and both the PVS and SAS are inhabited by fibroblasts and macrophages, as described below.

### Leptomeningeal Fibroblasts

In addition to vascular cells, fibroblasts are the main cell type found in the subarachnoid and perivascular spaces. These cells form fenestrated monolayers that cover blood vessels (pial sheath), the glia limitans (pia mater), and the inner layer of the arachnoid (Derk et al. 2021). Fibroblasts produce collagen, which provides structural integrity to both tissue and blood vessels. The primary function of pial and perivascular fibroblasts seems to be the secretion of collagen to reinforce the structural integrity of leptomeninges and blood vessels, rather than forming a fluid barrier.

Leptomeningeal fibroblasts are characterized by their squamous appearance, often displaying a disc-shaped nucleus (Figure 1d and e). These cells contain a low number of lysosomes and phagosomes and possess a primary cilium and fibropositors(Grubb 2022). In the MICrONS dataset, fibroblasts can also be found surrounding pial and parenchymal blood vessels, along with collagen fibrils that wrap around the blood vessels and traverse the PVS or SAS to adhere to the astroglial basement membrane, as I recently described(Grubb 2022).

Fibroblasts also contribute to the formation of the pia mater and arachnoid trabeculae (Figure 1d and e, Sup. Figure 1a and e), which span the SAS and suspend the brain, since the arachnoid mater adheres to the dura and the dura attaches to the skull(Haines 2018). These trabeculae are sometimes formed by fibroblast processes, cell bodies, or a combination of both (Figure 1e). Some macrophages also extend across the SAS (Sup. Figure 1a). However, the strength of the trabeculae is likely provided by large bundles of collagen fibers crossing the SAS and attaching to the inner arachnoid collagen matrix and to the subpial collagen matrix (Figure 1d and e, Sup. Figure 1a and e), rather than the cytoskeletons of fibroblast or macrophage processes.

The fibroblasts that form arachnoid trabeculae can also participate in the pial sheath, pia mater, or the inner arachnoid mater (Figure 1d). The primary cilia of fibroblasts are frequently in close contact with macrophages and can be observed invaginating the soma of macrophages with rounded cell bodies (Figure 1d and Sup. Figure 1b).

### Arachnoid Mater Structure

The arachnoid mater (Sup. Fig. 1a, Video 10) is composed an inner layer of inner arachnoid fibroblasts (Figure 1e.1) that attach to the inner arachnoid collagen matrix and form trabeculae that reach the subpial collagen matrix. Their morphology is indistinguishable from pia mater fibroblasts and they are likely the BFB2 cells recently identified (Pietilä et al. 2023). Above is a layer of reticular cells (Figure 1e.2), likely the same cells that have recently been named either SLYM or BFB3 fibroblasts(Møllgård et al. 2023; Orlin and Osen 1991; Pietilä et al. 2023). These are fenestrated cells that are loosely attached to eachother and to arachnoid barrier cells with adherence- and gap junctions (Sup. Fig. 1f and Sup. Figure 1c and Video 11) and are situated either in the collagen matrix or above it. They are recognizable by their large amount of rough endoplasmic reticulum and many large mitochondria (Sup. Fig. 1d) and often appear darker (more electron dense) than the rest of the cells in the arachnoid mater (Sup. Figure 1d). Reticular cells occasionally have more than one nucleus. Above is a layer of arachnoid barrier cells (Figure 1e.3). These cells form a tight barrier, are attached by long stretches adherence- and gap junctions (Sup. Fig. 1f) and no collagen fibers pass through them. They can be recognized by the slim and electron-dense intracellular space between them. Arachnoid barrier cells can be very large and contain up to 4 nuclei (Figure 1e.3 and Sup. Fig. 1e.3,4). The outer layer consists of loosely attached dural border cells that attach to arachnoid barrier cells with adherence- and gap junctions (Sup. Fig. 1f and Video 11). The cell bodies of the cells in the leptomeninges are dispersed, resulting in the entire arachnoid mater having a thickness approximately equal to two cell bodies. The collagen matrix traverses the layers (Sup. Figure 1a), forming 200-500 µm wide channels that run diagonally through the barrier layers, but crucially, they do not pass through the arachnoid barrier layer (Example, location: 146943, 89336, 17752). In areas where pial blood vessels are absent, the arachnoid mater appears continuous with the pia mater (Sup. Figure 1d). The adherens junctions between reticular cells or dural border cells with arachnoid barrier cells are likely composed of VE-cadherin, as this has been shown to be present in all the cell types(Mapunda et al. 2023). While the adherence junctions between arachnoid barrier cells can be either VE-cadherin or e-cadherin.

### Meningeal- and Perivascular macrophages: Function and Location

Macrophages play a critical role in the brain by consuming and digesting (phagocytosing) cellular metabolic waste, pathogens, and dead or damaged cells. Meningeal macrophages, characterized by their white phagosomes, multiple light-grey lysosomes (Sup. Figure 2a), and centrioles surrounded by the Golgi apparatus (Sup. Figure 2b, and not producing primary cilia), are numerous and evenly distributed in the subarachnoid space (SAS) atop pia mater fibroblasts (Figure 2a, Locations: tinyurl.com/345ybsch). Their tapered cell shape suggests active motility (Sup. Figure 2c, Video 2 and 7).

**Figure 2.**
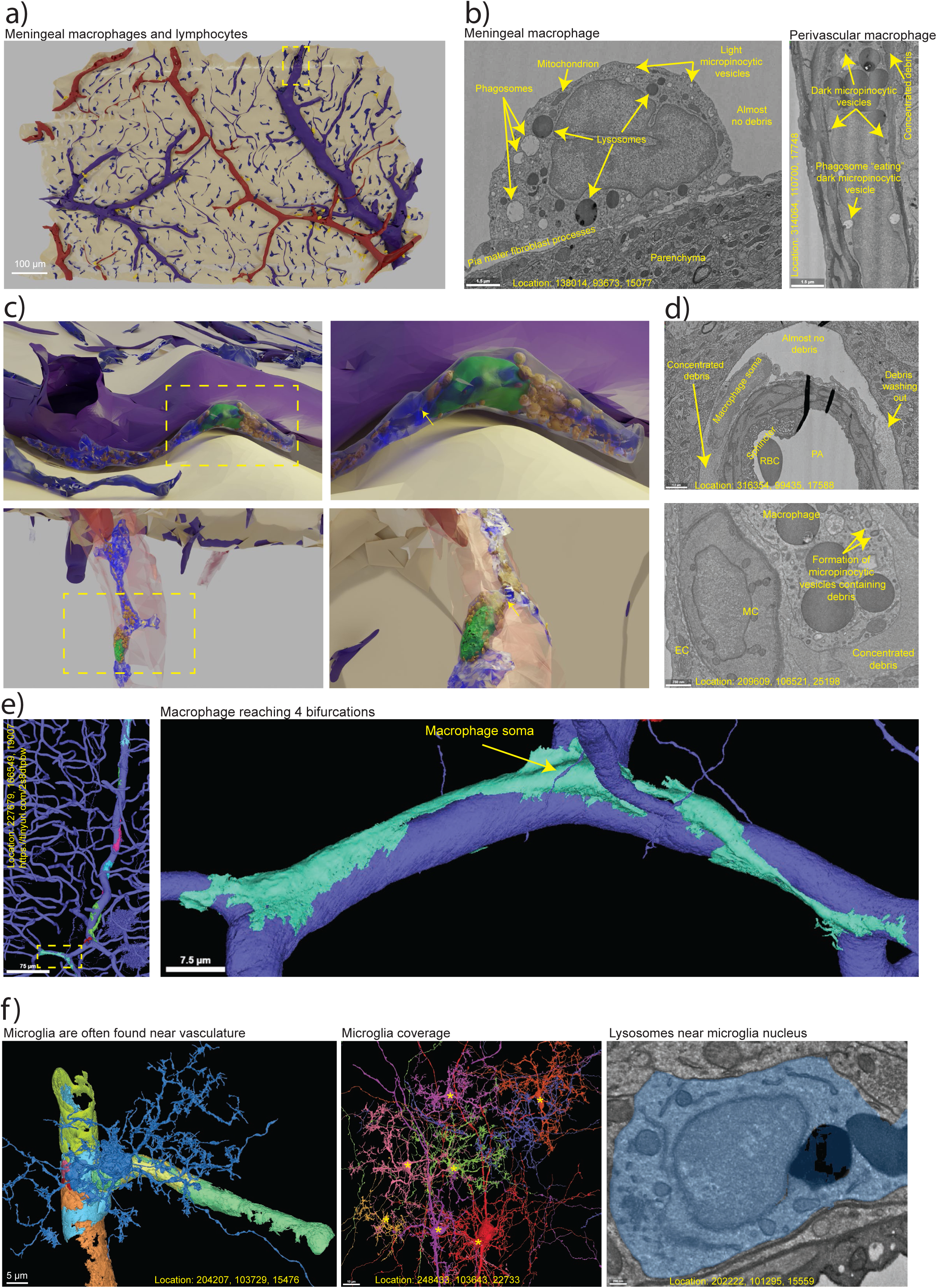
Macrophages and microglia ultrastructures. **a)** *Left panel:* Brain surface overview featuring meningeal macrophages (dark blue) and lymphocytes (yellow). The yellow dashed line indicates the location rendered in panel c. **b)** *Center panel:* Ultrastructure of a meningeal macrophage with electron-lucent micropinocytic vesicles. *Right panel:* Ultrastructure of a perivascular macrophage with electron-dense micropinocytic vesicles, matching the electron density of surrounding debris. **c)** *Top panels:* Rendered meningeal macrophage adjacent to a venule. *Lower panels:* Rendered perivascular macrophage in the PVS (hazy red) of a PA. Nucleus (green), lysosomes (orange) and centrioles (yellow). Yellow dashed lines show enlargement on the right and yellow arrows indicate centrioles locations. **d)** *Upper panel:* Ultrastructure reveals concentrated debris beneath a macrophage soma, seemingly obstructing debris washout. *lower panel:* Formation of micropinocytic vesicles containing debris with a concentration similar to that in the PVS. **e)** *Left panel:* 3D segmentation overview of a PA with 8 macrophages. *Right panel:* Example of a macrophage reaching 4 different bifurcations. **f)** *Left panel:* 3D segmentation of vascular-associated microglia (blue). *Central panel:* Volume coverage by 8 different microglia, with microglia cell bodies marked by yellow asterisks. *Right panel:* Ultrastructure of a typical microglia soma (blue) featuring two dark perinuclear lysosomes.

Perivascular macrophages are found on penetrating arterioles (PAs) and can reach up to the 3rd order capillaries (Figure 2c and e). For ascending venules, these macrophages are rare, found only near the brain surface and in a few cases (venules 5 and 6, Sup. Table 1). Metabolic waste debris is present in the deepest parts of perivascular spaces (PVSs), becoming diluted toward the SAS (Figure 2d and Sup. Figure 2d).

Occasionally, a fibroblast or macrophage cell body may occlude the PVS, causing debris to appear more concentrated (Figure 2d). Debris can also be found adhering to collagen fibrils (Sup. Figure 2d). Macrophagic micropinocytic vesicles contain debris (Figure 2b and Sup. Figure 2k), and vesicles can be observed being devoured by phagosomes (Figure 2b and Sup. Figure 2e), suggesting macrophages have a role in debris removal. Remnants of vesicle membranes can be found inside phagosomes (Sup. Figure 2e).

Phagosomes are often in close contact with lysosomes, which fuse together to enzymatically break down their contents (Sup. Figure 2f). Lysosomes also fuse with each other to produce larger lysosomes (Sup. Figure 2g). These larger lysosomes often contain fibrillar structures (Sup. Figure 2h) and may appear shredded by microtome slicing, particularly at electron-dense spots (Sup. Figure 2i). These spots are primarily found in meningeal macrophage lysosomes and less frequently in perivascular macrophages.

Perivascular fibroblasts often contain large heterolysosomes (Sup. Fig. 2i) which can be seen in the process of phagocytosis by macrophages (Sup. Fig. 2i), suggesting that macrophages aid fibroblasts in the removal of undigestable lysosome contents.

### Microglia Function and Identification

Microglia, originating from the same lineage as macrophages(Kierdorf et al. 2019), are found exclusively in the brain parenchyma and serve as an intrinsic immune defense. They continuously monitor the parenchyma for pathogens, plaques, and damaged or unnecessary synapses, phagocytosing these elements to prevent hyperexcitability in neurological diseases(Merlini et al. 2021). Microglia are distributed throughout the cortex, even near blood vessels (Figure 2d, tinyurl.com/3ej2kc56).

Like astrocytes, individual microglial cells cover distinct volumes of the parenchyma (Figure 2d, tinyurl.com/3n8n9a76). They can be identified by the presence of large electron-dense lysosomes near the nucleus and in larger processes (Figure 2f and Sup. Figure 2f). Phagosomes are present in both the cell body and processes and can be observed fusing with lysosomes (Sup. Figure 2f). Microglia are also seen pruning neuronal boutons or dendrites, which often contain phagosome-like vesicles—likely autophagosomes generated by oxidative stress(Talebi et al. 2022; Wendt et al. 2022).

### Lymphocyte Ultrastructural Characteristics

Lymphocytes are immune cells that regulate the activity of macrophages either by release of cytokines or by presentation of antigens via direct interaction. In the MICrONS dataset, lymphocytes are often found near vasculature (Figure 2a and Sup. Figure 2j), and in close contact with macrophages. Approximately half of the lymphocytes are located in the lower right corner of the brain surface, with none observed around parenchymal vessels.

Unlike fibroblasts, lymphocytes and macrophages lack primary cilia, and their centrioles are surrounded by the Golgi apparatus (Sup. Figure 2b). They often display multiple pseudopods (Sup. Figure 2j). Lymphocytes differ from macrophages in that they contain fewer and smaller lysosomes and phagosomes and have organelle-free areas in their cytosol (Sup. Figure 2j). There is one instance where a lymphocyte appears to be entering the arachnoid mater (Sup. Figure 2j).

### Arachnoid Villus and Macrophage Exocytosis

A tubular structure extending hundreds of micrometers in the subdural space, potentially an arachnoid villus, is observed in the MICrONS dataset (Figure 3, Sup. Figure 3a, tinyurl.com/ftce3f5s, Video 2 and 7). Internally, it is lined with dendritic cells featuring numerous abluminal pseudopods, large intermediate filament bundles(Kida et al. 1988) (Figure 3d), and sizable fibril-containing lysosomes. Externally, macrophages with large fibril-containing lysosomes sporadically cover the structure. The lumen contains a substance distinct from cerebrospinal fluid (CSF), appearing shredded by microtome slicing (Figure 3b, c and d).

**Figure 3.**
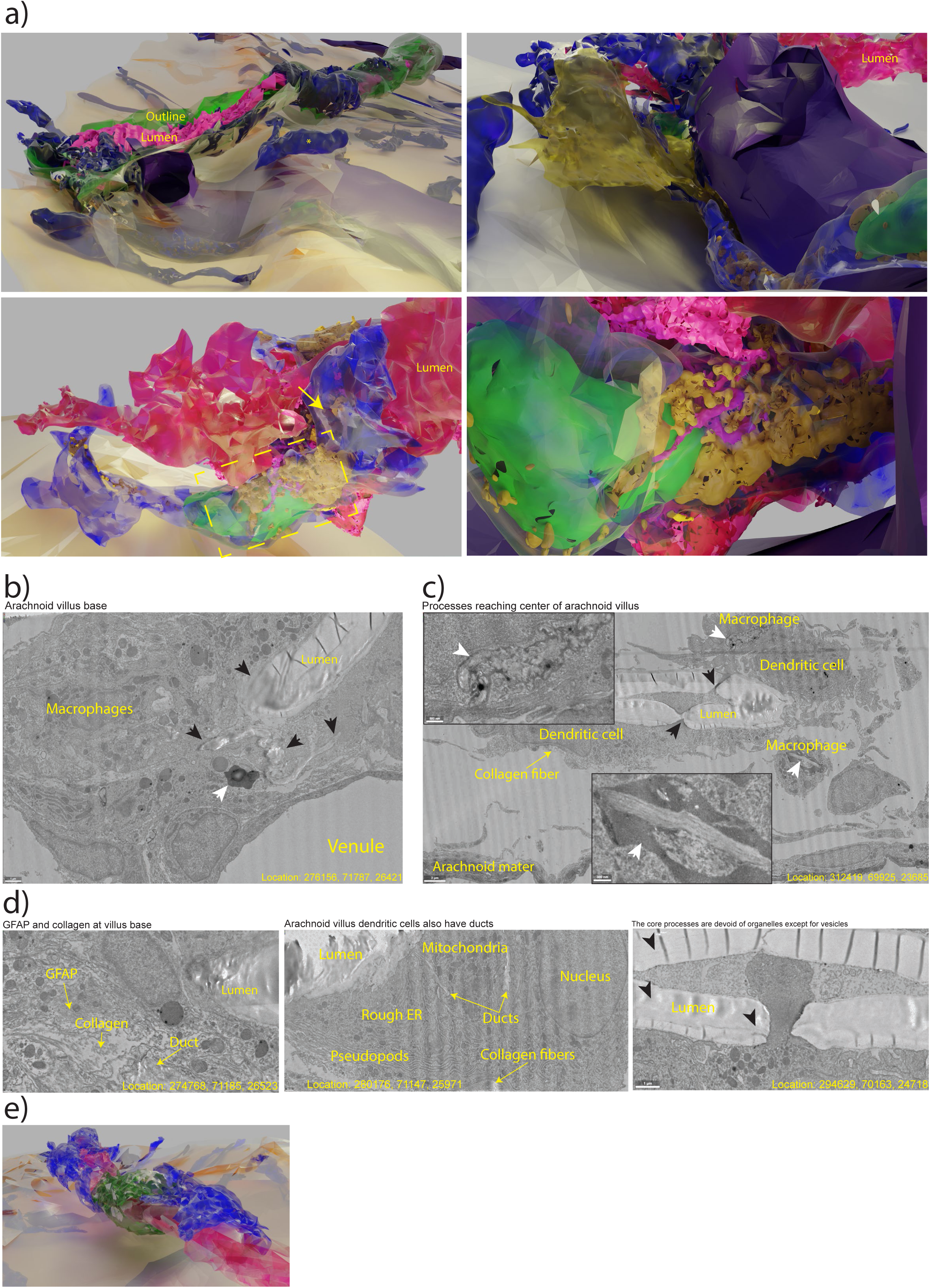
Arachnoid villus. **a)** *Upper left panel:* Meningeal macrophage location from Figure 2c with the arachnoid mater layer added, revealing an arachnoid villus crossing the subdural space towards a subdural venule. The villus outline is green, and the lumen is pink. The upper part of the villus base and the venule crossing the arachnoid mater appear severed, as they are at the dataset edge. Six macrophages are located at the villus base (left side of the image) beside the severed venule. Macrophages are distributed along the arachnoid villus stalk, and a subdural macrophage lies on the arachnoid mater (yellow asterisk). *Upper right panel:* View of the villus base from below the arachnoid mater (not included). Two arachnoid barrier cells (yellow) illustrate how macrophages move between them to access the villus base (see also Sup. Figure 3d and Video 9). *Lower left panel:* The villus base lumen with a single macrophage, showcasing the nucleus (green), lysosomes (orange), and centrioles (yellow). Yellow arrow indicates centriole location. *Lower left panel:* Close-up of the perinuclear region with exocytosed lysosomes forming ducts (pink, intracellular) that end in the villus lumen. Lysosome fusion is a segmentation error caused by poor EM picture alignment at the MICrONS dataset border. **b)** Ultrastructure of the arachnoid villus base displaying a shredded/folded substance in the villus lumen and ducts (black arrows). Villus base macrophages have large (Ø>2µm) lysosomes with lipofuscin or fibril inclusions (white arrow). **c)** Dendritic cell processes lining the villus lumen also cross the lumen (black arrows), forming a core in the arachnoid villus. Macrophages outside the arachnoid villus have many fibrils in their lysosomes and form elongated lysosomal inclusions resembling the villus-duct and lumen substance (white arrows). Insets: zoomed views of macrophage lysosomes. See also location: 334980, 69163, 22332. **d)** *Left panel:* Collagen fibers strengthen the villus structure, and GFAP from massive glia limitans astrocytes (Location: 270130, 72550, 26617) with large (Ø = 1-2 µm) GFAP bundles also reach the villus lumen (Location: 276906, 71055, 26272). *Center panel:* Dendritic cells lining the arachnoid villus lumen have 1-3 nuclei, extensive rough and smooth ER, and intracellular ducts leading to the villus lumen. *Right panel:* Core processes and most cytosol near the villus are devoid of organelles. Fibrillar patterns are visible in the villus lumen (black arrowheads). **e)** Rendered dendritic cell (green) and macrophages (blue) surrounding the arachnoid villus lumen (pink).

The structure’s base crosses the arachnoid mater alongside a large arachnoid-penetrating venule and extends towards a subdural venule, though it exits the dataset before they connect (Figure 3a). Six closely packed macrophages are located in the base complex, which is unusual given their even distribution in the SAS. These macrophages feature invaginations forming ducts that collect in the extracellular space and lead to the structure’s lumen (Figure 3a and b). The ducts also directly contact the endothelium of a nearby pial venule (Sup. Figure 3c).

Lysosomes in these macrophages are exceptionally large (Figure 3b) and occasionally resemble the extracellular ducts (Figure 3c), with contents also shredded by microtome slicing. This suggests ducts may form through lysosome exocytosis(Andrews 2000). Indeed, some lysosomes appear to be releasing their contents into the ducts (Sup. Figure 3b).

At the villus lumen’s center, cell processes of lining dendritic cells and macrophages form a core structure (Figure 3c and Video 3), which thickens with increasing distance from the villus base. Arachnoid barrier cells and fibroblasts externally line the structure towards the SAS (Figure 3a), and a meningeal macrophage is entering the arachnoid villus base between arachnoid barrier cells (Sup. Figure 3d). Large collagen fibers spiral around the arachnoid villus in the extracellular space, likely providing structural integrity (Figure 3c and d and Supp. Fig. 2d). Near the subdural venule, a macrophage is also observed crossing the inner arachnoid fibroblast layer and releasing lysosome content (Location: 336861, 67963, 24103 – putative lysosome exocytosis: 336838, 68337, 24175).

### Dendritic Cells in an Ectopic Lymphoid Aggregate

Dendritic cells are immune cells that are less common than macrophages and lymphocytes in the SAS. Both macrophages and dendritic cells differentiate from monocytes. Like macrophages, dendritic cells possess phagosomes and lysosomes. They are often multinucleated with multiple centrioles and primary cilia. The MICrONS dataset contains only a few dendritic cells, aside from those lining the arachnoid villus.

In one area at the brain surface’s lower right corner, an ectopic lymphoid aggregate comprising over 10 dendritic cells, several macrophages, and lymphocytes encapsulates very electron-dense particles (Figure 4a and b and Video 4). The exact identity of these particles is uncertain; however, based on their maximum size (∼50-80 nm), oval shapes, and surrounding immune cell density, they could be viruses. Since the brain is sliced into 40 nm sections, each particle may only appear in a single image. The electron-dense particles are also found in vesicles within the encapsulating dendritic cells (Sup. Figure 4a). Neutrophil extracellular trap (NET)-like structures, possibly restricting viral diffusion, are visible within the encapsulation (Figure 4b). One dendritic cell has 3 primary cilia (Sup. Figure 4b).

**Figure 4.**
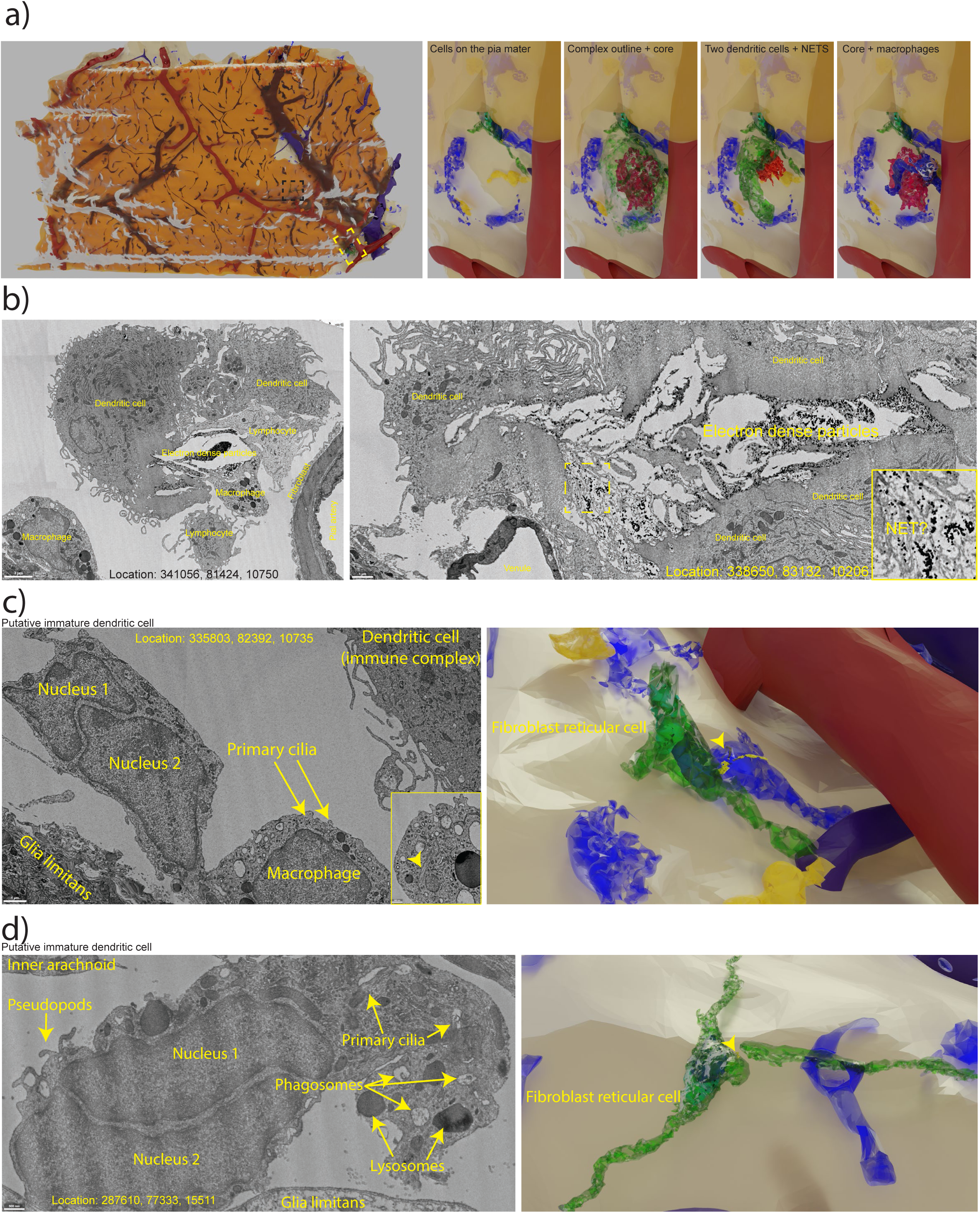
Ectopic lymphoid aggregate and dendritic cells. **a)** *Left panel:* Overview of immune cells beneath the arachnoid mater, highlighting the ectopic lymphoid aggregate location in the lower right corner (yellow dashed box) and a putative fibroblast reticular cell (black dashed box). *Right panels:* The ectopic lymphoid aggregate displayed in different combinations. The first panel from the left shows pial cells beneath the ectopic lymphoid aggregate, macrophages (blue), lymphocytes (yellow), and an fibroblast reticular cell (green). The second panel displays the entire ectopic lymphoid aggregate outline (hazy green) and the core (pink). The third panel features two dendritic cells within the ectopic lymphoid aggregate (green) and putative NETs (red). The fourth panel presents the core (pink) with macrophages (dark blue) in the ectopic lymphoid aggregate. **b)** *Left panel:* Ultrastructure of the ectopic lymphoid aggregate with electron-dense particles in the core surrounded by immune cells. *Right panel:* Different location displaying putative NETs. Yellow dashed box indicates enlarged inset in lower right corner. **c)** *Left panel:* Ultrastructure of a putative fibroblast reticular cell, also shown in Figure 4a, with two nuclei and two primary cilia. One cilium invaginates a nearby macrophage, and the inset in the lower right corner demonstrates the depth of the primary cilium within the macrophage. *Right panel:* Rendering of the putative fibroblast reticular cell (green) with primary cilia (yellow) invaginating the macrophage (blue). *Right panel:* Rendering of the putative fibroblast reticular cell (green) with primary cilia (yellow) invaginating the macrophage (blue). **d)** *Left panel:* Ultrastructure of a putative fibroblast reticular cell with two nuclei and two primary cilia. The cilia originate deep within the cell and only protrude slightly. *Right panel:* Rendering of the putative fibroblast reticular cell (green) with primary cilia (yellow, yellow arrowhead marks their location) invaginating the macrophage (blue). A dendritic cell process appears to have broken off, likely after fixation and before microtome slicing.

A cell with a segmented nucleus and multiple marginated electron-dense chromatin aggregates (Sup. Figure 4c), but few other organelles, is also encapsulated within the ectopic lymphoid aggregate. This cell could be an infected lymphocyte(Kiupel et al. 2001). A couple of fibroblast reticular cells, each with two nuclei and two primary cilia, are found in the SAS in the dataset’s lower right corner (Figure 4c and d). One putative dendritic cell’s primary cilium forms a deep invagination into a nearby macrophage (Figure 4c). Interestingly, one of the putative dendritic cells also contains very dilated smooth ER (Sup. Figure 4d), indicating high proteins synthesis and storage.

### Astrocytes and GFAP: Structure and Function

Astrocytes serve as a barrier between neurons and cerebrospinal fluid (CSF), with their endfeet surrounding blood vessels and forming the glia limitans at the brain surface. One notable ultrastructural feature of astrocytes is their dense bundles of glial fibrillary acidic protein (GFAP), which supports and strengthens them while maintaining their star shape. Though GFAP is a common astrocyte marker, it only labels 60% of them and is upregulated in reactive astrocytes(Walz and Lang 1998).

In the MICrONS dataset, GFAP bundles are easily identified in glia-limitans astrocytes (Figure 5a) and those on PAs and ascending venules in the cortex’s upper layers (Figure 5b), where there is a PVS. However, GFAP bundles are generally absent in the absence of a PVS (Figure 5f) and locating GFAP bundles in deeper cortical regions can be difficult. Glia-limitans astrocytes are characterized by extensive GFAP bundles, as shown by the uninterrupted bundles stretching from the pial surface to the endfeet, bypassing the nucleus, and ensheathing a layer 1 parenchymal venule (Figure 5a and Video 5).

**Figure 5.**
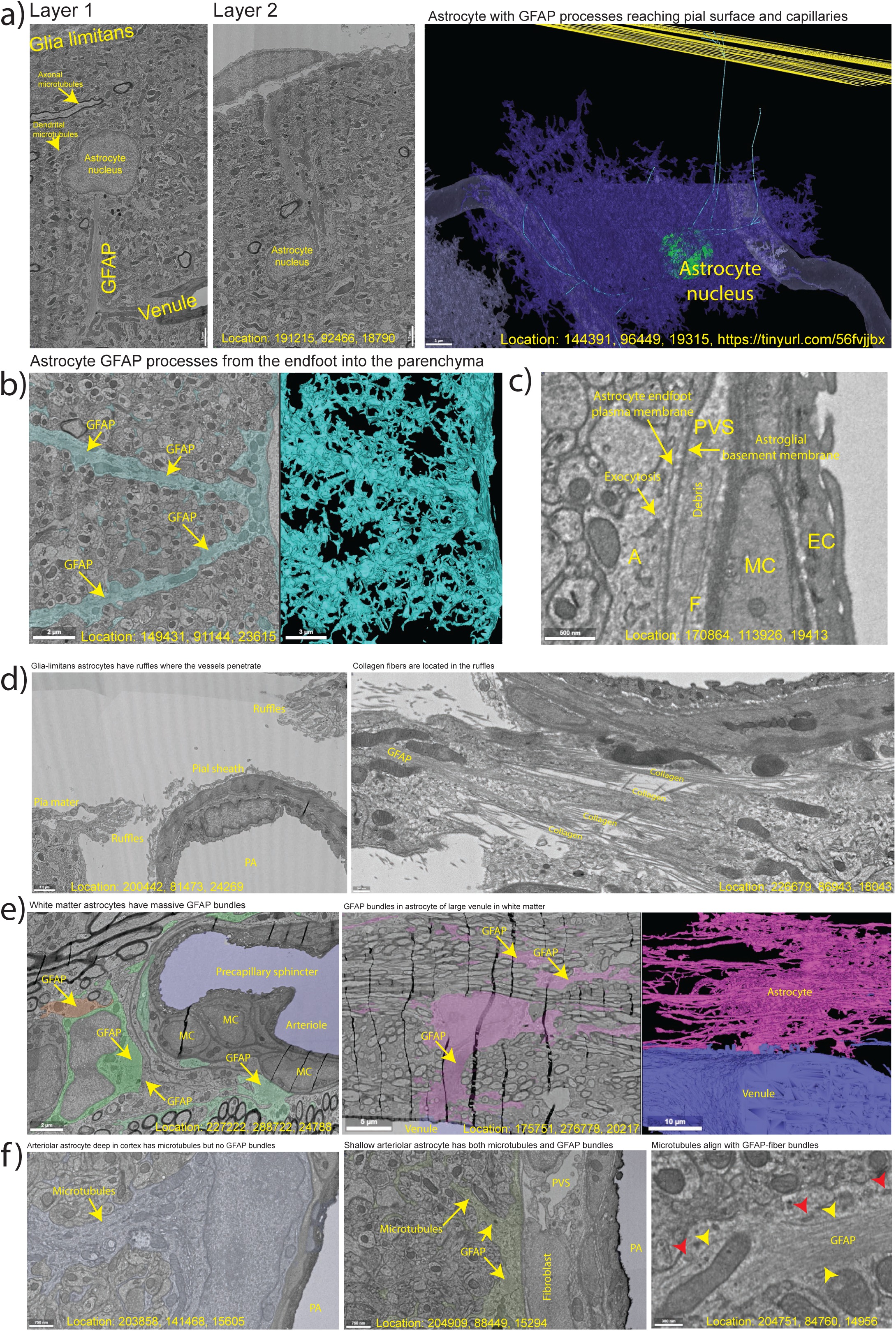
Astrocytes and GFAP. **a)** Ultrastructure of a glia limitans astrocyte (see Video 5) displays large GFAP bundles connecting the pial glia limitans with endfoot covering a parenchymal venule. *Left panel:* GFAP bundles reach the endfoot plasma membrane and can be distinguished from microtubules by differences in diameter. *Central panel:* GFAP bundle extending to the glia limitans. *Right panel:* Though the glia limitans is not segmented in the MICrONS dataset, GFAP bundles in an astrocyte (dark blue segmentation) can be traced (light blue lines) from the glia limitans at the brain surface (yellow lines) to two different capillaries (light blue segmentation) located near the astrocyte nucleus (green sphere). **b)** *Left panel:* Ultrastructure of a PA astrocyte (light blue) with processes radiating from the endfoot out into the parenchyma. GFAP bundles are indicated. *Right panel:* 3D segmentation of the astrocyte. **c)** Ultrastructure of an astrocytic endfoot (A) reveals exocytosis of a tubular vesicle into the PVS beneath the astroglial basement membrane. The CSF in the PVS contains metabolic waste debris. F = fibroblast process, MC = mural cell, EC = endothelial cell. **d)** *Left panel:* Ultrastructure of glia limitans around a PA exhibits ruffles in the astrocyte membrane and astroglial basement membrane. *Right panel:* GFAP bundles and collagen fibers support the ruffles from the inside and outside, respectively. **e)** *Left panel:* Ultrastructure of a white matter arteriole with a precapillary sphincter. The astrocytes (green, red and teal) contain massive GFAP bundles. *Center panel:* Ultrastructure of a white matter astrocyte (pink) with endfoot on a venule and GFAP bundles. *Right panel:* 3D segmentation of the astrocyte and venule. **f)** *Left panel:* Ultrastructure of an astrocyte surrounding a PA deep in the cortex, revealing the absence of GFAP bundles but presence of microtubules. *Center panel:* Ultrastructure of an astrocyte surrounding a PA shallow in the cortex, showing both GFAP bundles and microtubules in the processes. *Right panel:* Ultrastructure of an astrocyte process featuring GFAP bundles, microtubules (yellow arrowheads) at the GFAP periphery, and tubular electron-dense vesicles (red arrowheads) along the microtubules.

Impressively large GFAP bundles can have widths exceeding 500 nm and lengths over 20 µm. These massive bundles cause organelles, such as nuclei, to bend around them (Video 5). Despite their size, GFAP bundles contain no organelles, with only a few vesicle inclusions. However, mitochondria and long, tubular, electron-dense vesicles are observed along the GFAP bundles (Figure 5f). Some of these tubular vesicles can be seen emptying their content into the PVS (Figure 5c).

Glia-limitans astrocytes around large PAs display luminal membrane ruffles supported by collagen fibers, where the large GFAP bundles support the ruffles intracellularly (Figure 5d and Video 6). These ruffles increase the surface area of glia-limitans astrocytes, potentially enhancing water absorption. White matter astrocytes display massive GFAP bundles, unlike those in the deep cortical layers (Figure 5e). GFAP can be distinguished from microtubules based on the thickness of individual fibrils. Microtubules, nearly twice the size of GFAP superspirals(Margiotta and Bucci 2016; Tykhomyrov, Pavlova, and Nedzvetsky 2016), are often aligned with GFAP in astrocytes and can be found around GFAP bundle perimeters, along with tubular vesicles and mitochondria (Figure 5f).

## Discussion

The primary exit route for CSF and brain metabolic waste remains a topic of ongoing debate. The central question is whether metabolic waste exits the brain through the lymphatic system (Alves De Lima, Rustenhoven, and Kipnis 2020; Louveau et al. 2018; Ma et al. 2017) or via arachnoid villi or granulations(Shabo and Maxwell 1968; Turner 1958). The role of macrophages and arachnoid villi or granulations in metabolic waste removal may have been underestimated, as will be discussed in the following section.

### Arachnoid Mater Forms a Barrier

Here I show that the arachnoid mater consists of at least 4 different cell types, the dural border cells, the arachnoid barrier cells, reticular cells and inner arachnoid fibroblasts (Figure 1 and Sup. Fig. 1). These cells form a barrier that limits CSF to the SAS and any waste that is produced by the brain must somehow exit through this barrier. In the following, I am arguing that this waste is primarily filtered by macrophages before exiting the brain via specialized structures called arachnoid villi.

### Metabolic Waste Clearance in the Brain: Follow the Waste

Metabolic waste is produced by the activity of parenchymal cells, with some waste being cleared locally and much of it being released into the CSF(Iliff et al. 2012). In the parenchyma, metabolic waste encounters either microglia or astrocytes. Microglia may phagocytose and subsequently break down metabolic waste within their lysosomes, while astrocytes may play a role in transporting waste out of the parenchyma. As previously described(Grubb 2022), astrocytic endfeet overlap tightly, making a paracellular route between astrocytic endfeet seem unlikely for waste clearance. Instead, waste clearance may occur through vesicular transport via the microtubule network (Figure 5f) and exocytosis into either the PVS (Figure 5C) or, for glia limitans astrocytes, into the subpial space. Debris found in PVSs around PAs (Figure 2d, 5c and Sup. Figure 2k) likely represents metabolic waste that has not yet been cleared from the brain. From there, waste may be phagocytosed by perivascular macrophages (Figure 2 and Sup. Figure 2) or flow towards the SAS where numerous macrophages reside and where it may be removed by arachnoid granulations/villi or lymphatic drainage(Proulx 2021). The abundance of lysosomes and phagosomes in meningeal and perivascular macrophages suggests these cells are highly active phagocytes (Sup. Figure 2a). In ascending venules, the absence of PVSs and perivascular macrophages could imply that waste from astrocyte transcytosis is directly transcytosed into the venules by endothelial cells. Macrophages are evenly distributed across the pia mater, and the processes of meningeal macrophages extend along the brain surface or up to the arachnoid mater (Sup. Figure 1a and Video 8), with many pseudopodia providing a large surface for capturing floating waste. Perivascular pumping of waste towards the SAS could occur during deep-sleep stages(Fultz et al. 2019; Grubb and Lauritzen 2019; Mestre et al. 2018), when PVS volume fluctuates significantly(Bojarskaite et al. 2022). It has been suggested that perivascular and meningeal macrophages, in addition to immune surveillance, may play a role in filtering and draining the CNS(Kierdorf et al. 2019), with the density of perivascular macrophages being crucial for amyloid-β plaque clearance(Karam et al. 2022). This hypothesis is supported by the gradient in debris concentration towards the top of the brain and local accumulation of waste beneath obstructing fibroblast or macrophage cell bodies (Figure 2d). Additionally, perivascular macrophages surrounded by high concentrations of debris have dark micropinocytic vesicles, in contrast to meningeal macrophages with no surrounding debris (Figure 2d and Sup. Figure 2k).

### Macrophage Distribution and the Role of Fibroblasts in Waste Clearance

Although macrophages are numerous on the brain surface (Figure 2a), they rarely make contact with one another. This organization may be orchestrated by pial or inner-arachnoid fibroblasts, which possess primary cilia that extend to touch macrophages or form invaginations into macrophage somas. The exact role of fibroblast primary cilia, whether purely sensory or involving chemical or mechanical influences on macrophages, remains unclear. However, the even distribution of macrophages could optimize the waste-filtering effect of phagocytosing macrophages. Perivascular fibroblasts generate collagen fibrils that span the PVS(Grubb 2022). These fibrils connect mural cells with astrocytic endfeet and may explain why the astrocyte-endfoot tube follows PA constrictions during sleep(Bojarskaite et al. 2022), causing perivascular pumping(Mestre et al. 2018). In line with this, perivascular macrophages have been shown to regulate arterial motion(Drieu et al. 2022), likely by breaking down perivascular collagen fibrils.

### Macrophage Migration and Fibrillary Protein Clearance at Arachnoid Villi

The mechanism by which macrophages handle fibrillary proteins, which may be difficult or impossible to degrade and thus accumulate in lysosomes, remains unclear. However, studies have shown that conditions within lysosomes facilitate the formation of fibrillar structures(Bayati et al. 2022; Schützmann et al. 2021). An MRI study demonstrated that macrophages migrate along parenchymal blood vessels(Mori et al. 2014). I propose that macrophages move to the pial surface to release their fibrillary lysosome contents. In the MICrONS dataset, I present a structure next to a venule that penetrates the arachnoid mater, which may be a murine arachnoid villus, where macrophages gather and release their lysosome content. The higher incidence of macrophage lysosomes containing fibrils near the putative arachnoid villus (Figure 3c, Sup. Figure 2h and 3d and f) compared with perivascular macrophages (Sup. Figure 2a), supports the idea that macrophages migrate to arachnoid villi to exocytose their fibril-containing lysosomes. This suggests that arachnoid villi could serve as an exit route for fibrillar metabolic debris, such as α-synuclein or β-amyloid.

### Arachnoid Villi Lumen Substance and Lysosome Exocytosis

The arachnoid villus contains a substance that appears similar to what is seen in TEM images of arachnoid granulations in the venous sinus subendothelial space of dogs(Andres 1967), sheep(Jayatilaka 1965), primates and humans(Shabo and Maxwell 1968; Turner 1958) (Sup. Figure 3e). Turner described the substance as elastic tissue with wavy or occasionally coiled appearance (Turner 1958). Jayatilaka observed that electron-dense bodies (lysosomes) of macrophages in the arachnoid granulations of sheep contain regularly oriented parallel fibrils (Jayatilaka 1965) (Sup. Figure 3f). I propose that, due to differences in brain size, the mouse arachnoid villi lumen corresponds to the arachnoid-villus subendothelial space in larger mammals. Moreover, the mouse arachnoid villi lumen substance is likely a mixture of exocytosed lysosomes and fibrillary proteins.

### Exit Routes for Metabolic Waste

Additionally, debris may also exit through the cribriform plate in the nasal cavity, where CSF drains directly from the SAS into nasal lymphatics(Kida, Pantazis, and Weller 1993). Recent studies using fluorescent tracers have suggested that some dural lymphatic vessels, aside from those at the cribriform plate, may serve as exit routes for CSF(Alves De Lima et al. 2020; Louveau et al. 2018; Ma et al. 2017; Proulx 2021), but it remains unclear how macromolecules might enter dural lymph vessels that are located on the far side of the arachnoid CSF barrier(Brinker et al. 2014), and unfortunately, the dura mater is not present in the MICrONS dataset. Identifying the primary exit route for metabolic waste may be complicated by the breakdown or inhibition of fluorescent tracers in macrophages. Since PEGylation protects against phagocytosis by macrophages, the use of PEGylated tracers may be biased towards the lymphatic route, bypassing arachnoid villi(Ma et al. 2017).

### Microglia-Exocytosis in Alzheimer’s

In accordance with the macrophage-exocytosis-in-arachnoid-villi theory, it is plausible that in diseases such as Alzheimer’s, microglial lysosomes accumulate fibrils, and plaques could represent local deposits where microglia release their fibrillary lysosome content into the extracellular space through exocytosis. This is supported by the observation that the amount of fibrillary material in microglia increases with their proximity to plaques(El Hajj et al. 2019). Similarly, I have found that macrophages near the arachnoid villus contain a higher quantity of lysosomal fibrillary material compared to other pial or perivascular macrophages. In line with these findings, microglia depletion has been shown to reduce plaque formation but increase amyloid accumulation around blood vessels, leading to cerebral amyloid angiopathy(Spangenberg et al. 2019).

### Pathologies Overwhelming Waste Clearance

Pathologies could potentially overwhelm the brain’s waste clearance system. For example, in subarachnoid hemorrhage, physical blockage caused by blood clotting and/or intense immune responses may disrupt the normal filtration of debris by macrophages and hinder the release of waste through exocytosis into arachnoid villi. This disruption could result in an accumulation of harmful aggregation-prone proteins, which may ultimately contribute to the development of dementia(Corraini et al. 2017).

A recent preprint has found that during chronic infection with Trypanosoma brucei, the meninges acquire lymphoid-tissue properties and autoreactive B-lymphocytes(Alcala et al. 2023). They find a structure (their sup. fig. 5) closely resembling the arachnoid villus structure shown in this study. The lymphoid-like structure also consists of dendritic cells (CD21/CD35^+^) lining the lumen, and when infected it attracts B- and T-lymphocytes (B220^+^/CD3d^+^). Importantly, this structure also exists in the uninfected mouse, but without the lymphocytes. It will be interesting to see whether macrophages adhere to this structure.

### Meningeal Immune Responses

In addition to waste clearance, meningeal macrophages and dendritic cells play a critical role in containing and removing pathogens such as bacteria and viruses. Infections can lead to meningitis, which can be life-threatening in bacterial cases, but generally less severe for viral infections. Interestingly, the MICrONS dataset features an instance of meningeal dendritic cells and macrophages encapsulating a potential viral infection in the SAS (Figure 4). Studying this example in greater detail may provide valuable insights into immune reactions within the brain.

### Microglia Ensuring Blood-Brain Barrier Integrity

A recent study discovered that microglia directly associate with pericytes(Morris et al. 2023). Utilizing the MICrONS dataset, the authors suggested a direct contact between microglial and pericyte cell bodies through an astrocytic endfoot fenestration. However, the provided example only shows the astrocytic endfoot becoming very thin, and it is unclear if there is direct contact. In another instance, a microglial cell was found to plug several gaps between astrocytic endfeet (Locations: 257544, 179071, 25585 and 257195, 181050, 25575 and 254260, 179303, 25874 and 254497, 179035, 25928), consistent with a previous study by Misje Mathiisen et al. (2010) demonstrating that microglia fill in holes in astrocytic endfoot coverage(Misje Mathiisen et al. 2010). Additionally, Haruwaka et al. (2019) reported that microglia exert a dual effect on systemic inflammation. Initially, they help maintain blood-brain barrier integrity by physically contacting endothelial cells past the astrocyte endfeet. However, at later stages, microglia phagocytose the astrocyte endfeet. These findings emphasize the complexity of microglia-pericyte interactions and their implications for the blood-brain barrier and astrocytic endfoot coverage(Haruwaka et al. 2019).

### Astrocytic Endfeet and GFAP: Implications for Water Regulation and Metabolite Clearance

Astrocytic endfeet are connected by gap junctions, similar to endothelial cells(Simard et al. 2003). However, it is unclear whether they also form tight junctions. In a previous study(Grubb 2022), I observed that astrocyte somas form endfeet on both arterioles and venules, with significant overlap. This observation questions the notion that paracellular diffusion between astrocytic endfeet is the primary route for brain water and solute exchange and instead suggests that astrocytic endfeet may act as a barrier. In contrast, astrocytes could be involved in cortical water uptake and solute movement into or out of the parenchyma through a transcellular route(Rasmussen et al. 2022).

GFAP expression in astrocytes suggests a role beyond providing structural integrity. GFAP is an intermediate filament protein that influences the localization of aquaporin 4 channel (AQP4), Na^+^, K^+^, 2Cl^−^, and water cotransporter (NKCC1), and other proteins involved in water and ion regulation(Wang et al. 2016). Interestingly, GFAP is also highly expressed in kidney glomeruli podocytes(Buniatian et al. 2002), which are responsible for the glomerular filtration barrier. This suggests that GFAP may play a role in water filtration.

GFAP may also be involved in astrocytic transcellular water movement by decreasing resistance to intracellular flow or by providing capillary forces. This process would require an intracellular-osmolarity gradient from one endfoot to another or from an endfoot to the astrocyte parenchymal processes. GFAP upregulation in ischemic penumbra or around fibrotic scars could be an adaptation to counter swelling(Wang et al. 2016). Supporting this idea, the lack of GFAP is associated with hydrocephalus and dysmyelination in the white matter of aging mice(Liedtke et al. 1996). Overall, the study of astrocytic endfeet and GFAP highlights their potential implications in water and ion regulation, metabolite clearance, and the maintenance of brain homeostasis.

## Conclusion

I propose that brain waste clearance primarily involves a pathway comprising microglia, astrocytes, perivascular macrophages, perivascular pumping, meningeal macrophages, and arachnoid villi. While CSF may exit the brain via the lymphatic system, metabolic debris is likely filtered by macrophages (or microglia) through phagocytosis and enzymatic degradation in lysosomes, followed by lysosome exocytosis into the arachnoid villi. This "gliarachnoid" system could become overwhelmed by pathological conditions that stress or obstruct the involved cells, potentially leading to dementia.

Furthermore, I propose that waste and water exchange between the parenchymal interstitial fluid and CSF follows a transcellular route through astrocytes. GFAP may play a central role in astrocyte water homeostasis. Understanding the complex interactions between these cell types and GFAP in the context of waste clearance and water regulation could provide valuable insights into maintaining brain homeostasis and preventing neurodegenerative diseases.

## Limitations

The locations where tissue disruption occurred after fixation but before microtome slicing can be identified by the disrupted plasma membranes of the cells constituting the tissue. Tissue fixation is known to shrink the extracellular space and possibly the perivascular space (PVS) as well. However, the extreme pial perivascular-space shrinkage observed in Mestre et al., where the pial artery collapsed and there was likely significant tissue swelling, is not evident in the MICrONS dataset(Mestre et al. 2018). In this dataset, both pial arteries and venules maintain a slightly oval shape, and the subarachnoid space (SAS) appears to be intact.

It is possible that the arachnoid villus structure is not an arachnoid villus but rather an early stage of bone regrowth (osteogenesis). Osteogenesis is known to occur in mice with cranial windows, and a durotomy has also been performed for this mouse(Consortium et al. 2021). Furthermore, the intense electron density of the putative virus particles raises questions about their true identity. An alternative explanation could be that the particles are calcium phosphate. The virus-like particles are also found inside vesicles of a cell with three nuclei and three primary cilia, which could be an early osteoblast (Sup. Figure 4a). It is known that macrophages can fuse and become osteoclasts, and that calcium phosphate can mediate that process(Wang et al. 2021). This evidence supports the alternative hypothesis that the putative arachnoid villus is arachnoid osteogenesis initiated due to craniotomy and durotomy.

However, bone regrowth in cranial windows grows inward as a sheet from the skull fragments at the periphery at a maximum rate of 250 µm/week(Roome and Kuhn 2014), rather than as long tubular structures. Since the diameter of the MICrONS dataset craniotomy is 4 mm and the mice were fixed 8 days after surgery (1 day for recovery, 6 days for imaging, and overnight transport to the Allen Institute), the expected regrowth would limit the craniotomy to a diameter of 3.5 mm. This is more than double the length (1.4 mm) and more than triple the width (0.87 mm) of the rectangular MICrONS dataset, making it unlikely that the edges of the dataset are near the bone regrowth zone.

## Methods

### Public volume electron microscopy dataset

This work is based on the MICrONS dataset provided by the Allen institute(Consortium et al. 2021), as described previously(Bonney et al. 2022; Grubb 2022). The dataset comprises an approximately 1 mm^3^ chunk of the mouse (male P75-87) visual cortex, including parts of the primary visual cortex and higher visual areas for all cortical layers except extremes of L1. The dimensions are (*in vivo*) 1.3 mm mediolateral, 0.87 mm anterior-posterior and 0.82 mm radial. The dataset also includes the arachnoid mater; however, the dura mater is not present. The resolution was initially ∼4 nm but was downsampled to 8 nm/pixel, and the slice thickness was set at 40 nm. The dataset consists of two subvolumes, with one containing 65% of the total volume and the other containing 35% of the total volume. A vascular annotation for both volumes can be found at: tinyurl.com/48vf8pdd. Movies were created by capturing multiple screenshots using a macro created in iCue, a gamer keyboard software. The screenshots were then cropped, stacked, and converted to ".avi" format using ImageJ software.

### 3D rendering

Vascular annotations were exported from Neuroglancer as JSON code and trimmed in Notepad++ using a regex code with help from ChatGPT to obtain XYZ coordinates. These coordinates were opened in Meshlab and a mesh was added. The meshes were opened in Blender and after cleaning of the meshes, light and camera settings were set before rendering. For all 3D renderings the scalebars are only approximate. The brain model used in Video 9 is from https://scalablebrainatlas.incf.org/, the skull model is from http://www.mouseimaging.ca the mouse model is https://hackday.yahoo.co.jp/.

## Acknowledgements

I would like to express my sincere gratitude to the Translational Neuroscience lab at the Department of Neuroscience, University of Copenhagen, for their invaluable scientific discussions and insights during my six years at the institution. I am also grateful to the Novo Nordisk Foundation, the Lundbeck Foundation, the Danish Medical Research Council, the Alice Brenaa Fondation, and a Nordea Foundation Grant to the Center for Healthy Aging for providing financial support through grants for Professor Martin Lauritzen.

## Supplementary figure texts

**Supplementary Figure 1**

**a)** *Left panel:* Ultrastructure of the arachnoid mater, a trabecular fibroblast, and a pial-sheath fibroblast with collagen fibers (black arrowheads) visible between the layers of arachnoid barrier cells. *Central panel:* Macrophages extend from the pia to the arachnoid, resembling a trabecula. *Right panel:* Collagen fibers span the distance between the arachnoid- and the pia mater. **b)** An example of an inner arachnoid fibroblast with a primary cilium reaching through the SAS to make contact with a meningeal macrophage (Location: 143768, 89303, 17140). **c)** Example of three adherence junctions between an arachnoid barrier cell and a reticular cell. **d)** Example of a location without any blood vessels, where the arachnoid mater is continuous with the pia mater. **e)** Supplementary views for Figure 1e.1) Sideview of the inner arachnoid fibroblast and collagen matrix. 2) Topview showing the small reticular cell with only 1 nucleus. 3) The large arachnoid barrier cell shown without any other cell covering. 4) The small arachnoid barrier cell shown without any other cell covering. **f)** Segmentation of the adherens junctions within a 10 µm^3^ cube. Left panel: Segmentation shown as colored lines on top of orthogonal views of the EM data. Right panel: Adherens junctions and gap junctions (GJ) of arachnoid barrier cells (ABC) between themselves, with dural border cells (DBC) or with reticular cells (RC).

**Supplementary Figure 2**

**a)** Periarteriolar macrophage (light green segmentation) exhibiting a high number of phagosomes and relatively small lysosomes. **b)** Example of Golgi apparatus (yellow arrow) encircling the centrioles in a macrophage, a feature also observed in lymphocytes. **c)** Line annotations (green) illustrating the approximate direction of movement for meningeal and perivascular macrophages as a vector from the nuclei (X) to the centrioles (no X) – see tinyurl.com/2uwyyjyp. **d)** Debris in the PVS around a PA adhering to collagen fibrils. **e)** *Upper panel:* Vesicle-membrane remnants (black arrowhead) found inside phagosomes. *Lower panel:* Example of a phagosome engulfing a micropinocytic vesicle (black arrowhead) and a lysosome that has internalized phagosomes (white arrowhead). **f)** *Left panel:* Lysosomes observed fusing with phagosomes in microglia. *Right panel:* Microglia in the process of phagocytosing pre-synapses containing large vacuoles. **g)** Example of lysosomes fusing in a macrophage. Note how some of the more electron-dense lysosome content from one lysosome appears in the other lysosome where they fuse (yellow arrow). **h)** Example of fibrillary structures in a large meningeal macrophage lysosome. **i)** Example of microtome shredding of electron-dense spots in macrophage lysosomes. **j)** *Upper panel:* A lymphocyte with few organelles and pseudopods. *Lower panel:* Lymphocyte moving across the inner arachnoid fibroblast layer. **k)** Example of a macrophage in the PVS of a PA with dark debris in the CSF (black arrow) and in the micropinocytic vesicles (yellow arrow). **i)** Example of heterolysosomes in perivascular fibroblasts location: 261296, 147378, 25443. **j)** Example of a macrophage phagocytosing a perivascular fibroblast heterolysosome, location: 163662, 221448, 18458. Other examples are locations: 160922, 223483, 18402 and 159651, 222964, 18294 and 160073, 222680, 18323 and 261360, 145398, 25480 and 266335, 177892, 25224.

**Supplementary Figure 3**

**a)** *Upper left panel:* Neuroglancer 3D annotation of a portion of the arachnoid villus structure. The villus base is situated next to a venule, and is directed towards another venule which it contacts. **b)** *Left panel:* Ultrastructural example of a macrophage lysosome at the arachnoid villus base fusing with an extracellular duct. *Right panel:* Dendritic cell lysosome at the arachnoid villus base fusing with the villus lumen. Observe the dark substance along the edge of the lumen. **c)** An extracellular duct is seen in close proximity to venule’s endothelium. The yellow dashed box marks the location of the enlarged inset on the right side, which demonstrates that the duct penetrates the arachnoid barrier cells, the inner arachnoid fibroblast layer, the pial sheath and endothelial basement membrane and forms invaginations into the venular endothelium. **d)** Meningeal macrophage crossing the arachnoid barrier cell layer to enter the arachnoid villus base. The yellow arrow marks the leading macrophage process. **e)** Arachnoid granulations in the literature in dog, sheep, monkey and humans reveal a substance (yellow arrows) in the subendothelial space that is similar to the content of the mouse arachnoid villus lumen. **f)** Example of a macrophage lysosome with fibrillary content in an arachnoid granulation of a sheep.

**Supplementary Figure 4**

**a)** A dendritic cell within the ectopic lymphoid aggregate contains vesicles with electron-dense particles of similar size to the putative viruses. **b)** Another dendritic cell in the ectopic lymphoid aggregate possesses 3 nuclei and 3 primary cilia (yellow arrowheads). **c)** An encapsulated apoptotic cell features a segmented nucleus with condensed chromatin, possibly indicating an infected lymphocyte. **d)** The putative fibroblast reticular cell in Figure 4d exhibits severely dilated smooth ER, likely indicative of high protein production.

## Supplementary video texts

**Video 1**

Vascular segmentations in the MICrONS dataset, along with annotations of the glia limitans (brown), the arachnoid mater (yellow), pial vasculature (arterioles = red, venules = blue) and PVSs (arteriolar PVS = light pink, venular PVS = light blue).

**Video 2**

Annotations of meningeal macrophages (dark blue) and of macrophage nucleus and centrioles (green). The green lines mark the shortest distance from the nucleus to the centrioles. The upper limit of the white matter is also annotated (white). The video zooms in to show the location of the arachnoid villus base (outline = orange, lumen = pink) and rotates around the base to show the extracellular ducts (pink). For a 3D rendered version of the villus base, see video 9.

**Video 3**

Annotations of a dendritic cell (teal) lining the lumen of the arachnoid villus, showcasing dendritic cell processes reaching the villus core through ultrastructural images.

**Video 4**

Annotations of an ectopic lymphoid aggregate (virus capture) in the SAS. Lymphocytes (gold) are also shown, and they are numerous close to the ectopic lymphoid aggregate (outline = orange, core = cyan). Ultrastructural images are shown to illustrate how the structure of the ectopic lymphoid aggregate, NETS (red), dendritic cell (yellow), macrophages (dark blue, purple or pink), lymphocytes and fibroblast reticular cell (golden).

**Video 5**

Ultrastructure of glia limitans ruffles surrounding a pial artery (PA), supported by GFAP (intermediate filaments) and collagen. Notice how close the GFAP filaments get to the glia limitans membrane.

**Video 6**

Ultrastructure of the ruffles of glia limitans around a PA. GFAP (intermediate filaments) and collagen support the ruffles.

**Video 7**

View a rendered video illustrating the location and relative size of the MICrONS dataset in a mouse brain, using 3D models from Thingiverse.com (see methods section for exact sources) and a bedding texture from https://cloud.blender.org/p/textures/.

**Video 8**

Rendered video summarizing the study’s main findings, including immune cells, ectopic lymphoid aggregate, fibroblasts, macrophages, dendritic cells and arachnoid villus.

**Video 9**

Rendered video demonstrating how macrophages with dense lysosomes pass through arachnoid barrier cells to reach the arachnoid villus base, where they exocytose lysosomes into the villus lumen via ducts.

**Video 10**

Rendered video showing the layers of the leptomeninges with the arachnoid mater.

**Video 11**

Video showing the annotations of adherence junctions in the arachnoid mater.

---

**Supplementary Table 1:**
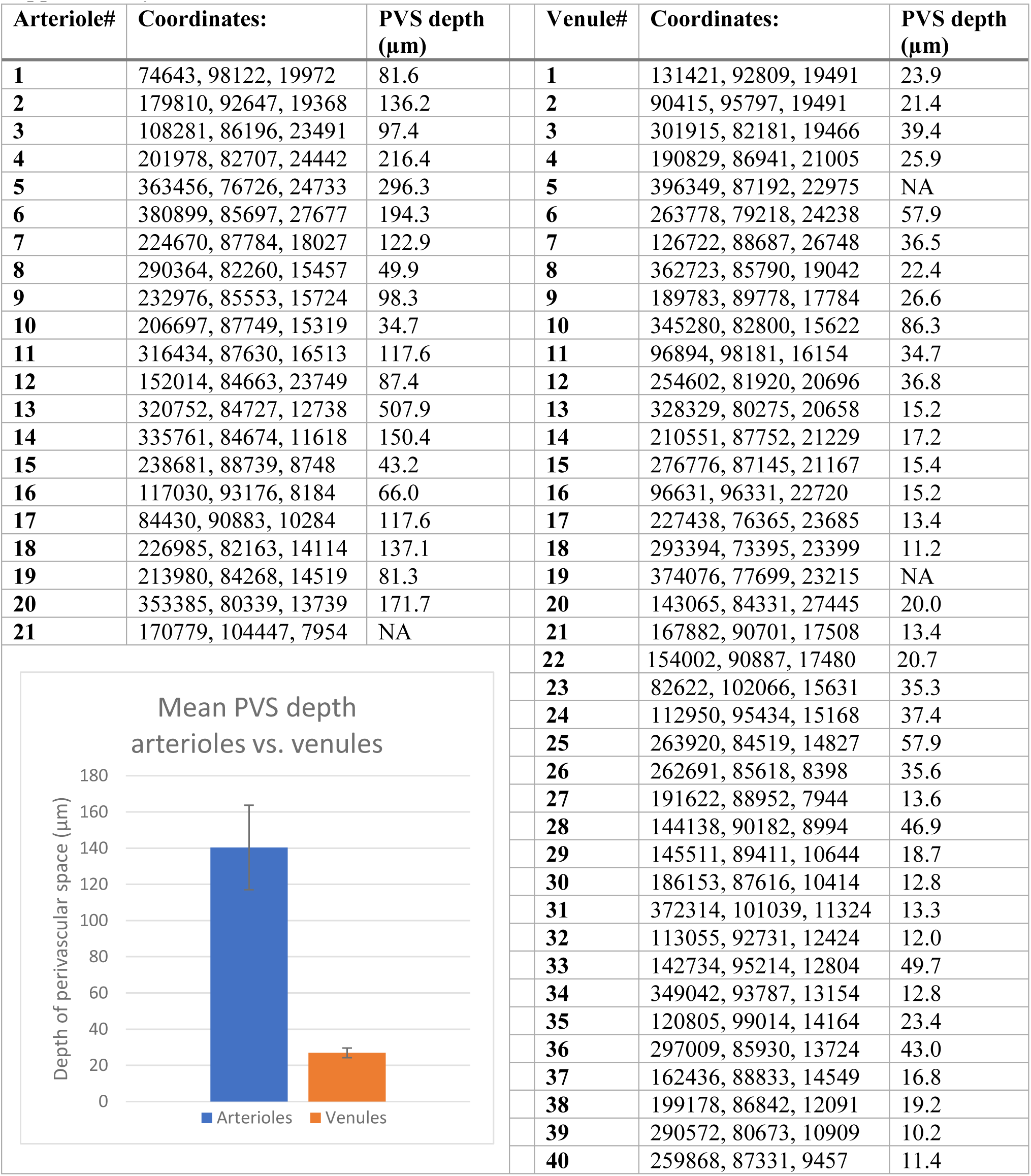

