## Supplementary figures and images for "Ultrastructure of the brain waste-clearance pathway"

### Sup. Fig. 1

#### Supplementary Figure 1

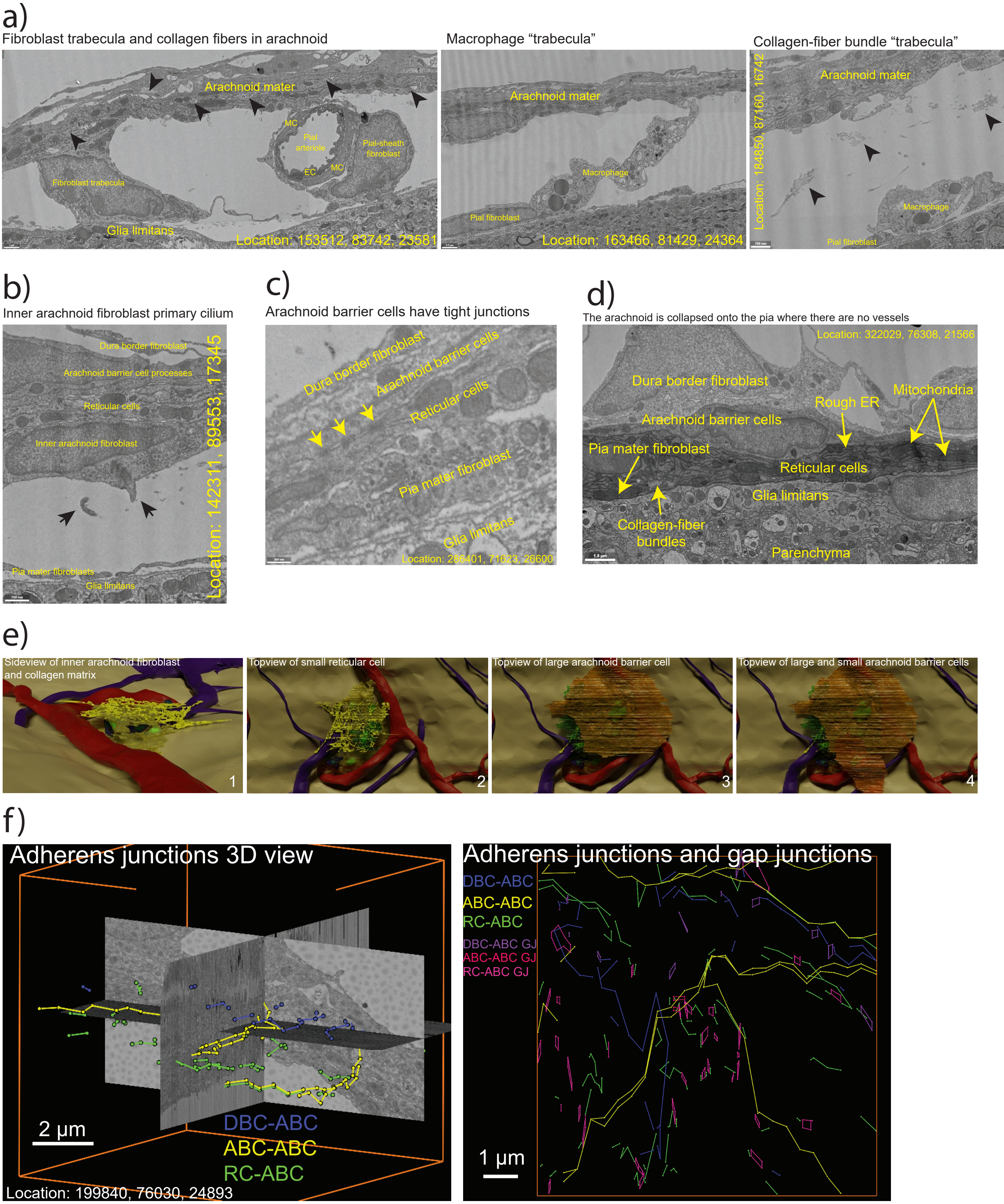

### Sup. Fig. 2

#### Supplementary Figure 2

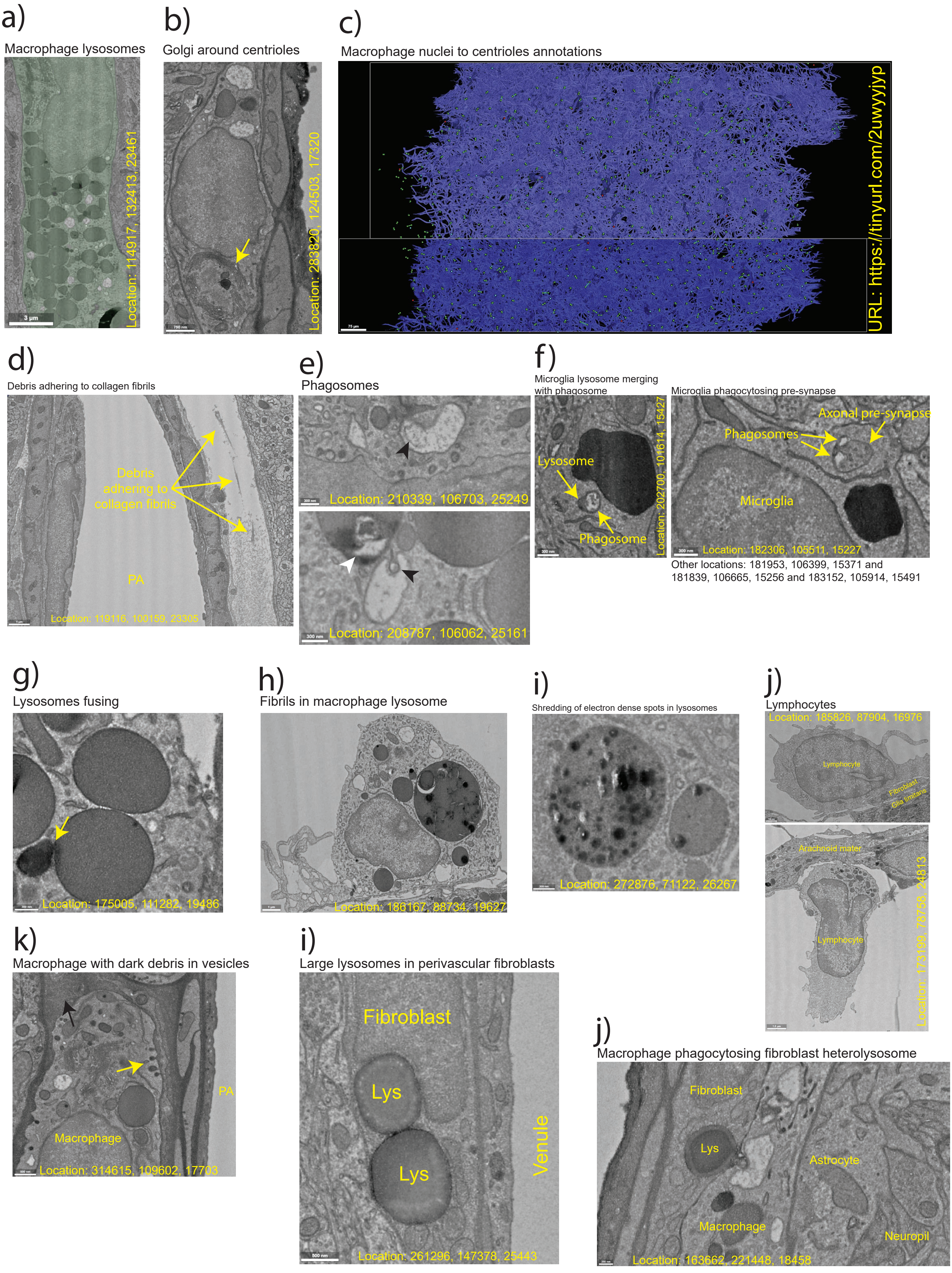
