## Supplementary material for "Ultrastructure of the brain waste-clearance pathway": Sup. Fig. 3

### Supplementary Figure 3

a)

Arachnoid villus crossing from one venule to another

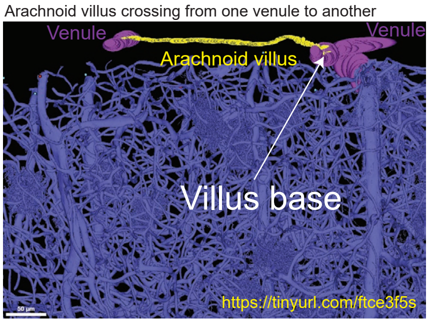


b)

Macrophage lysosome fusing with extracellular duct

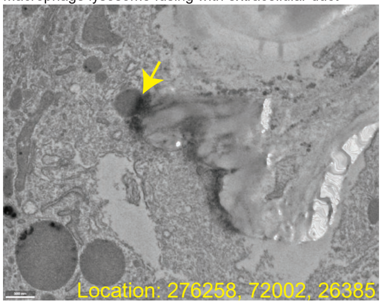


Dendritic cell lysosome fusing with lumen

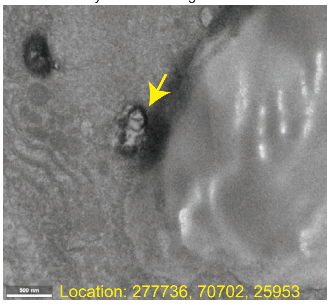


c)

Duct invaginating a venular endothelial cell

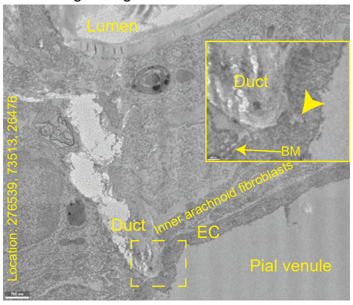


d)

Macrophage moving in between fibroblast processes at arachnoid villus base

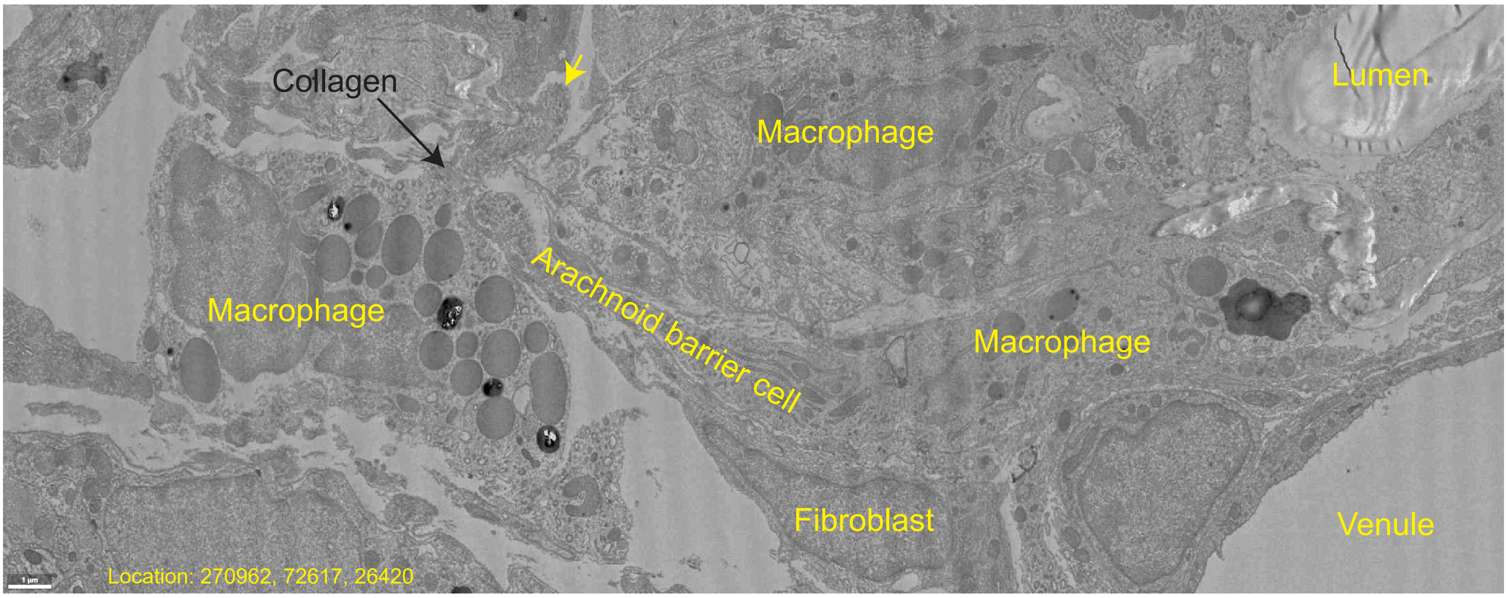


e)

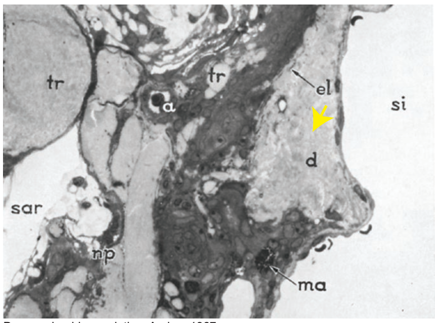


Dog arachnoid granulation, Andres 1967

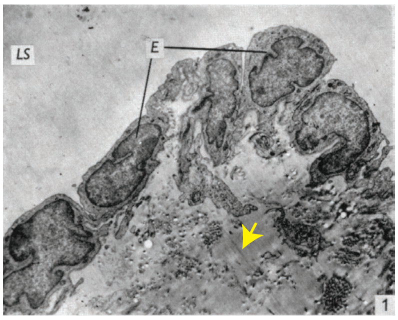


Sheep arachnoid granulation, Jayatilaka 1965

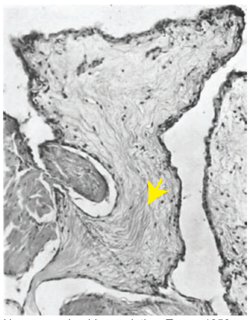

Human arachnoid granulation, Turner 1958

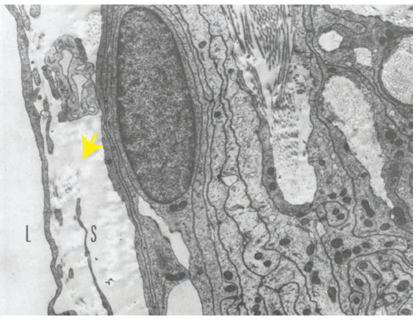


Monkey arachnoid granulation, Shabo and Maxwell 1968

f)

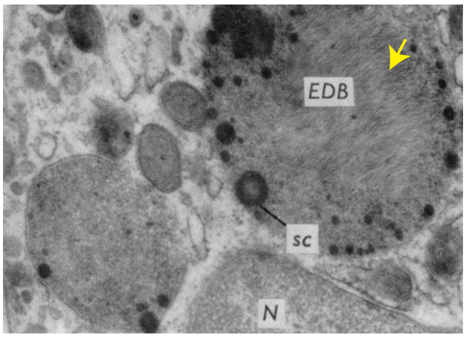


Macrophage lysosomes in a sheep arachnoid granulation, Jayatilaka 1965
