## Supplementary material for "Ultrastructure of the brain waste-clearance pathway": Sup. Fig. 4

### Supplementary Figure 4

a)

Dendritic cell with electron-dense particles in vesicles

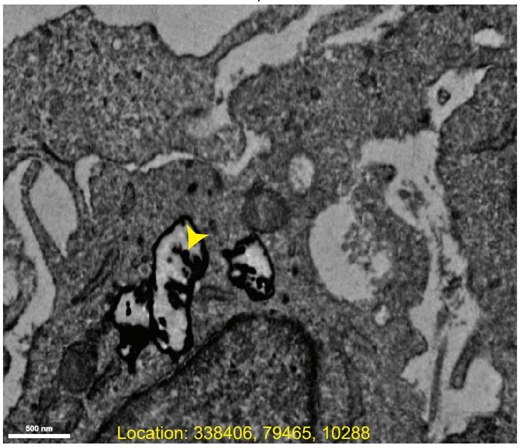


b)

Three primary cilia of the dendritic cell

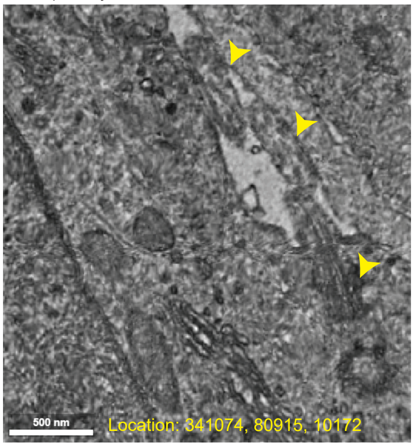


c)

Aggregated chromatin in encapsulated lymphocyte

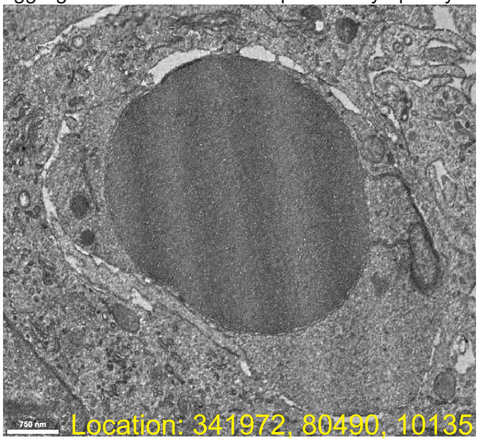


d)

Dilated smooth ER in dendritic cell

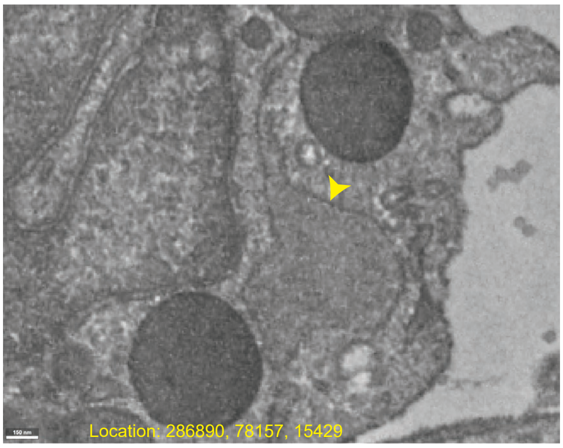
